# Chikungunya virus outbreak in the Amazon region: replacement of the Asian genotype by an ECSA lineage?

**DOI:** 10.1101/492595

**Authors:** F. G. Naveca, I. Claro, M. Giovanetti, J. G. Jesus, J. Xavier, F. C. M. Iani, V. A. do Nascimento, V. C. de Souza, P. P. Silveira, J. Lourenço, M. Santillana, M. U. G. Kraemer, J. Quick, S. C. Hill, J. Thézé, R. D. O. Carvalho, V. Azevedo, F. C. S. Salles, M. R. T. Nunes, P. S. Lemos, D. S. Candido, G. C. Pereira, M. A. A. Oliveira, C. A. R. Meneses, R. M. Maito, C. R. S. B. Cunha, D. P. S. Campos, M. C. Castilho, T. C. S. Siqueira, T. M. Terra, C. F. C. de Albuquerque, L. N. da Cruz, A. L. Abreu, D. V. Martins, D. S. M. V. Simoes, R. S. Aguiar, S. L. B. Luz, N. Loman, O. G. Pybus, E. C. Sabino, O. Okumoto, L. C. J. Alcantara, N. R. Faria

## Abstract

**Background:** Since its first detection in the Caribbean in late 2013, chikungunya virus (CHIKV) has affected 51 countries in the Americas. The CHIKV epidemic in the Americas was caused by the CHIKV-Asian genotype. In August 2014, local transmission of the CHIKV-Asian genotype was detected in the Brazilian Amazon region. However, a distinct lineage, the CHIKV-East-Central-South-America (ECSA)-genotype, was detected nearly simultaneously in Feira de Santana, Bahia state, northeast Brazil. The genomic diversity and the dynamics of CHIKV in the Brazilian Amazon region remains poorly understood despite its importance to better understand the epidemiological spread and public health impact of CHIKV in the country.

**Methodology/Principal Findings:** We report a large CHIKV outbreak (5,928 notified cases between August 2014 and August 2018) in Boa vista municipality, capital city of Roraima’s state, located in the Brazilian Amazon region. In just 48 hours, we generated 20 novel CHIKV-ECSA genomes from the Brazilian Amazon region using MinION portable genome sequencing. Phylogenetic analyses revealed that despite an early introduction of the Asian genotype in 2015 in Roraima, the large CHIKV outbreak in 2017 in Boa Vista was caused by an ECSA-lineage most likely introduced from northeastern Brazil.

Epidemiological analyses suggest a basic reproductive number of R_0_ of 1.66, which translates in an estimated 39 (95% CI: 36 to 45) % of Roraima’s population infected with CHIKV-ECSA. Finally, we find a strong association between Google search activity and the local laboratory-confirmed CHIKV cases in Roraima.

**Conclusions/Significance:** This study highlights the potential of combining traditional surveillance with portable genome sequencing technologies and digital epidemiology to inform public health surveillance in the Amazon region. Our data reveal a large CHIKV-ECSA outbreak in Boa Vista, limited potential for future CHIKV outbreaks, and indicate a replacement of the Asian genotype by the ECSA genotype in the Amazon region.

**Author Summary:** Until the end of 2017, Brazil notified the highest number of infections caused by chikungunya virus (CHIKV) in the Americas. We investigated a large CHIKV outbreak in Boa vista municipality in the Brazilian Amazon region. Rapid portable genome sequencing of 20 novel isolates and subsequent genetic analysis revealed that ECSA lineage was introduced from northeastern Brazil to Roraima around July 2016. Epidemiological analyses suggest a basic reproductive number of R_0_ of 1.66, which suggests that approximately 39% of Roraima’s population was infected with CHIKV-ECSA. Given the dominance of the CHIKV-Asian genotype in the Americas, our data highlights the rapid spread of a less understood and poorly characterized CHIKV-ECSA genotype in Brazil. Investigations on potential associations between public health impact of CHIKV and genetic diversity of circulating strains are warranted to better evaluate its impact in Brazil and beyond.

## Introduction

In August 2014, local transmission of chikungunya virus (CHIKV) was detected in Brazil for the first time, with cases being reported nearly simultaneously in Oiapoque (Amapá state, north Brazil) and Feira de Santana (Bahia state, northeast Brazil), two municipalities separated by >2000 km distance. Genetic analysis confirmed the co-circulation of distinct virus lineages in Brazil: the Asian genotype (CHIKV-Asian) was introduced to Oiapoque possibly from neighbouring French Guiana, while the East-Central-South-African genotype (CHIKV-ECSA) was introduced to Feira de Santana from a traveller returning from Angola [1].

Since 2014 and until the end of September 2018, a total of 697,564 CHIKV cases have been notified in Brazil (including 94,672 laboratory-confirmed cases). This is the largest number recorded in any of the 51 countries or territories reporting local CHIKV transmission in the Americas [2]. The virus has been circulating in the Americas since 2013 where approximately 260 million people live in areas at-risk of transmission [2-4]. Similar to the recent Zika virus epidemic [5], the rapid spread of CHIKV in the Americas, including in Brazil, results from several factors, including the establishment and abundance of competent *Aedes* spp. vectors, lack of population immunity, and increased mobility of vectors and humans between regions reporting current presence of the virus [6].

Chikungunya virus is an enveloped, non-segmented, single-stranded positive polarity RNA alphavirus that is a member of the *Togaviridae* family and is transmitted predominately by the *Aedes aegypti* and *Aedes albopictus* vectors, which are widespread in Brazil [7]. There are four main genotypes: (i) the West African genotype is maintained in an enzootic cycle in Africa, (ii) the Asian genotype, which is endemic in Asia, (iii) the East-Central-South-African genotype, endemic to Africa, and (iv) the Indian Ocean Lineage (IOL) genotype, an epidemic lineage that emerged from the ECSA genotype around 2004 and swept through the Indian Ocean region causing a series of explosive outbreaks [8].

The first symptoms of CHIKV infection are a rapid increase in temperature (>38.9°C), followed by severe, often debilitating polyarthralgia. Serological data from La Reunion, Philippines and the Indian Ocean island of Mayotte suggest that 75-97% of persons infected with CHIKV develop symptomatic infections [9]. Seroprevalence data from Brazil suggests that 45.7 to 57.1% Riachão do Jacuípe and of Feira de Santana, both located in Bahia state, were exposed to CHIKV in 2015, with a total of 32.7% to 41.2% of the population reporting symptoms [10].

Throughout Asia and the Americas, chikungunya virus outbreaks have been associated with unique clinical features [11], including long-lasting symptoms [12], and high mortality resulting from complications associated with CHIKV infection [13, 14]. In Brazil, a striking proportion of 68.1 to 75% of the population with positive serological results reporting symptoms contracted a chronic form of the disease [12, 15]. However, the epidemiological features, genomic diversity, and transmission dynamics of recent CHIKV outbreaks in this country remain poorly understood. Inferences that are based only on clinical-epidemiological notifications are complicated by underreporting of cases by the national reporting system [16], mostly due to the co-circulation and co-infection with viruses that cause overlapping symptoms, such as Zika and dengue viruses [17-19]. Moreover, CHIKV serological tests may cross-react with other alphaviruses, such as Mayaro virus, that circulate in the north and centre-west regions of Brazil [20, 21]. In this context, it is challenging to use only clinical-epidemiological and serological data to evaluate the true extent of the disease. Moreover, accurate incidence data is critical to forecast and provide prediction of the course of epidemics [22].

Until the end of 2016, 83.3% of the cases in Brazil were reported in northeast region of the country [23]. However, in 2017, Roraima state, located in the Amazon basin in the north of Brazil, reported its first large CHIKV outbreak. Roraima is the northernmost state of Brazil, lies in the Amazon basin, borders Venezuela and French Guiana to the north, and Amazonas and Pará states to the south, and its equatorial climate favours year round transmission of mosquito-borne viruses [24]. Within Brazil’s northern states, Roraima has been implicated as a stepping-stone to virus introductions from other Latin American regions, such as dengue [25], and yellow fever virus in the past [26]. Moreover, the Amazon region has recently been highlighted as a region with high transmission potential of vector-borne diseases [4] and, more generally, a region with high potential for virus zoonoses and emergence [27].

Due to its connectivity and potential impact on global epidemiology of vector-borne and zoonotic virus from the Amazon basin, it is important to improve genomic pathogen surveillance in Roraima. By August 2018, the public health laboratory of Boa Vista (capital city of Roraima state) had reported 5,928 CHIKV cases, 3,795 of which were laboratory-confirmed. Here we a use combination of on-site portable virus genome sequencing, and epidemiological analysis of case count and web search data to describe the circulation, genetic diversity, epidemic potential and attack rates of a large CHIKV outbreak in Boa Vista.

## Methods

### Connectivity in study area

Roraima is the northernmost of Brazil’s 27 federal units (**Figure 1a**) and has an estimated population of 450,479, of whom 284,313 live in the capital city of Boa Vista (ibge.gov.br/). Despite being Brazil’s least populated federal unit, Roraima is one of the best-connected Brazilian states in the Amazon basin [28]. Within Brazil, Roraima is connected to Amazonas state in the south via the road BR-174. This road also connects Roraima’s capital city, Boa Vista, to the states of Bolivar and Amazonas in Venezuela in the north. Further, the road BR-401 links Boa Vista to Guyana in the east. There are four daily flights connecting Boa Vista with Brasília, capital of Brazil, as well as six weekly flights to Manaus, the capital city of Amazonas state and the biggest city in the north of the country, with connecting daily nonstop flights to all other Brazilian states/regions and international destinations, including important international airport hubs in Panamá City and Miami, USA. There are also less-commonly used seasonal fluvial networks that connect Boa Vista and Manaus via the Amazonas river.

**Figure 1.**
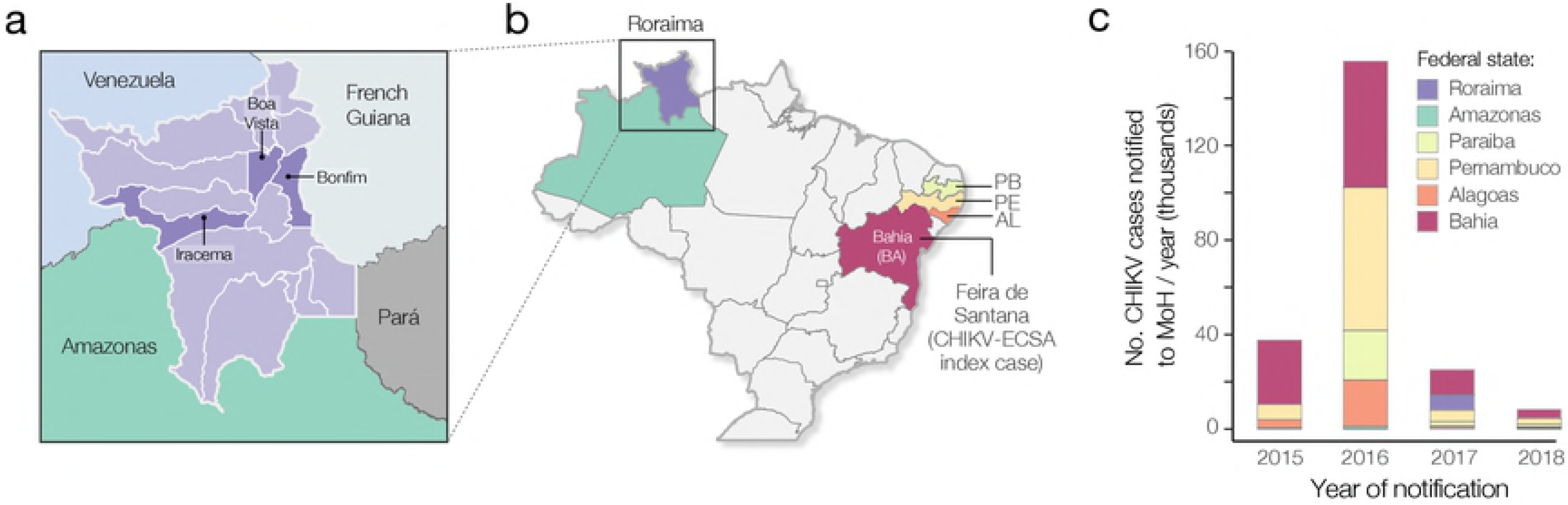
Context of this study. A. Map showing municipalities of Roraima state, including Boa Vista, bordering countries (Venezuela and French Guiana) and bordering Brazilian federal states (Amazonas and Pará). B. Map of Brazilian states, showing the states from which CHIKV sequence data in this study was analysed (Bahia, Alagoas, Pernambuco, Paraíba, Amazonas and Roraima). C. Barplot showing the annual number of notified CHIKV cases in selected states of Brazil (data obtained from the Brazilian Ministry of Health). Map was made with Natural Earth. Free vector and raster map data at naturalearthdata.com.

### Chikungunya virus case count time series

The Roraima State Central Laboratory (LACEN-RR) is responsible for the differential diagnosis of suspected arbovirus cases presenting to Roraima’s public health units. Between Jan 2014 and September 2018, LACEN-RR notified 5,928 CHIKV cases in Boa Vista alone, 3,795 of these laboratory-confirmed, to the National Reportable Disease Information System (SINAN). Case count time series are available from Github (http://github.com/arbospread). We follow the Brazilian Ministry of Health’s guidelines and define a notified CHIKV case as a suspected case characterized by (i) acute onset of fever >38.5°C, (ii) severe arthralgia and/or arthritis not explained by other medical conditions, and (iii) residing or having visited epidemic areas within 15 days before onset of symptoms. A laboratory-confirmed case is a suspected case confirmed by laboratory methods such as (i) virus isolation in cell culture, (ii) detection of viral RNA, (iii) detection of virus-specific IgM antibodies in a single serum sample collected in the acute or convalescent stage of infection; or (iv) a four-fold rise of IgG titres in samples collected during the acute phase, in comparison with a sample collected in the convalescent period.

### Nucleic acid isolation and RT-qPCR

Residual anonymized clinical diagnostic samples were sent to Instituto Leônidas e Maria Deane, FIOCRUZ Manaus, Amazonas, Brazil, for molecular diagnostics as part of the ZiBRA-2 project. The ZiBRA-2 project was approved by the Pan American Health Organization Ethics Review Committee (PAHOERC) n° PAHO-2016-08-0029. Total RNA extraction was performed with QIAmp Viral RNA Mini kit (Qiagen), following manufacturer’s recommendations. Samples were first tested using a multiplexed qRT-PCR protocol against CHIKV, dengue virus (DENV1-4), yellow fever virus, Zika virus, Oropouche virus and Mayaro virus [29]. All qRT-PCR results were corroborated using a second protocol [30]; comparable Ct values were obtained with the two protocols. CHIKV positive samples tested negative for all other arboviruses tested. Samples were selected for sequencing based on Ct-value <30 (to maximize genome coverage of clinical samples by nanopore sequencing [31]), and based on the availability of epidemiological metadata, such as date of onset of symptoms, date of sample collection, gender, municipality of residence, and symptoms (**Table 1**). We included 13 samples from Roraima state plus 5 additional samples from patients visiting the LACEN-Amazonas in Manaus, under the auspices of the ZiBRA project (http://www.zibraproject.org/). All samples were processed in accordance with the terms of Resolution 510/2016 of CONEP (National Ethical Committee for Research, Brazilian Ministry of Health).

**Table 1.** Epidemiological data for virus isolates from Roraima (RR) and Amazonas (AM). CT=cycle threshold, *d*=days from onset of symptoms to sample collection. Corresponding sequencing statistics are available in **Table S1**. Isolates were collected around 2.3 (range: 0 – 5) days after onset of symptoms. Acc. Number = GenBank accession number.

| Isolate | State,<br>Municipality | Acc.<br>Number | Ct RT-qPCR | Coverage<br>(%) | Age,<br>Sex | Collection<br>date | *d* |
| --- | --- | --- | --- | --- | --- | --- | --- |
| AMA290 | AM, Manaus | MK121891 | NA | 90.2 | 76, F | 15/07/2015 | 5 |
| AMA291 | AM, Manaus | MK121892 | NA | 80.7 | 48, F | 15/07/2015 | 4 |
| AMA292 | AM, Manaus | MK121893 | NA | 90.2 | 50, M | 15/07/2015 | 0 |
| AMA293 | AM, Manaus | MK121894 | NA | 84.4 | 42, M | 31/01/2016 | 4 |
| AMA294 | RR, Boa Vista | MK134712 | NA | 90.2 | 45, F | 01/12/2014 | 2 |
| AMA295 | RR, Unknown | MK134713 | NA | 90.2 | 9, F | 11/11/2014 | 1 |
| AMA74 | AM, Manaus | MK121895 | 15 | 90.2 | 32, F | 20/03/2017 | 2 |
| AMA346 | RR, Boa Vista | MK121896 | 13.7 | 90.2 | 30, F | 03/03/2017 | 1 |
| AMA350 | RR, Bonfim | MK121897 | 27.15 | 54.7 | 32, F | 20/02/2017 | 1 |
| AMA352 | RR, Boa Vista | MK121898 | 17.33 | 88.6 | 3, F | 22/02/2017 | 1 |
| AMA354 | RR, Boa Vista | MK121899 | 23.36 | 86.9 | 19, F | 17/03/2017 | 1 |
| AMA362 | RR, Iracema | MK121900 | 18.63 | 88.6 | 31, F | 17/03/2017 | 1 |
| AMA364 | RR, Boa Vista | MK121901 | 25.93 | 83.3 | 19, F | 17/03/2017 | 2 |
| AMA366 | RR, Boa Vista | MK121902 | 19.87 | 90.0 | 36, F | 17/03/2017 | 2 |
| AMA368 | RR, Boa Vista | MK121903 | 25.91 | 93.1 | 26, F | 15/03/2017 | 2 |
| AMA369 | RR, Boa Vista | MK121904 | 21.55 | 95.6 | 52, M | 02/03/2017 | 3 |
| AMA374 | RR, Boa Vista | MK121905 | 27.41 | 71.4 | 64, F | 02/03/2017 | 4 |
| AMA379 | RR, Boa Vista | MK121906 | 17.5 | 96.1 | 38, F | 27/02/2017 | 4 |
| AMA381 | RR, Boa Vista | MK121907 | 16.66 | 97.7 | 31, F | 27/02/2017 | 4 |
| AMA382 | RR, Boa Vista | MK121908 | 14.58 | 76.6 | 30, F | 05/03/2017 | 1 |

### Complete genome MinION nanopore sequencing

Between the 1^st^ and 7^th^ June 2017, we attempted sequencing at Instituto Leônidas e Maria Deane, FIOCRUZ Manaus on all selected samples with Ct-value <30. We used an Oxford Nanopore MinION device with protocol chemistry R9.4, as previously described [32]. Sequencing statistics can be found in **Table S1 (Julien)**. In brief, we employed a protocol with cDNA synthesis using random primers followed by strain-specific multiplex PCR [32]. Extracted RNA was converted to cDNA using the Protoscript II First Strand cDNA synthesis Kit (New England Biolabs, Hitchin, UK) and random hexamer priming. CHIKV genome amplification by multiplex PCR was attempted using the CHIKAsianECSA primer scheme and 35 cycles of PCR using Q5 High-Fidelity DNA polymerase (NEB) as described in [32]. PCR products were cleaned up using AmpureXP purification beads (Beckman Coulter, High Wycombe, UK) and quantified using fluorimetry with the Qubit dsDNA High Sensitivity assay on the Qubit 3.0 instrument (Life Technologies). PCR products for samples yielding sufficient material were barcoded and pooled in an equimolar fashion using the Native Barcoding Kit (Oxford Nanopore Technologies, Oxford, UK). Sequencing libraries were generated from the barcoded products using the Genomic DNA Sequencing Kit SQK-MAP007/SQK-LSK108 (Oxford Nanopore Technologies). Libraries were loaded onto a R9/R9.4 flow cell and sequencing data were collected for up to 48 hr. Consensus genome sequences were produced by alignment of two-direction reads to a CHIKV virus reference genome (GenBank Accession number: N11602) as previously described in [32]. Positions with ≥ 20× genome coverage were used to produce consensus alleles, while regions with lower coverage, and those in primer-binding regions were masked with N characters. Validation of the sequencing protocol was previously performed in [32].

### Collation of CHIKV-ECSA complete genome datasets

Genotyping was first conducted using the phylogenetic arbovirus subtyping tool available at http://www.krisp.org.za/tools.php. Complete and near complete sequences were retrieved from GenBank on June 2017 [33]. Two complete or near-complete CHIKV genome datasets were generated. Dataset 1 included ECSA-PreAm (ECSA sampled outside the Americas) and ECSA-Br (ECSA sequences sampled in the Americas) sequences. This dataset contained 36 complete genomes from the ECSA genotype, including 7 from East and Central Africa (HM045823 from Angola 1962; HM045784 from Central African Republic 1984; HM045812 from Uganda 1982; KY038947 from Central African Republic 1983; HM045793 from Central African Republic 1986; HM045822 from Central African Republic 1978; and KY038946 from Central African Republic 1975). Dataset 1 also included 29 sequences from Brazil, including the new 18 genomes reported here from the ECSA lineage and 3 genomes from the outbreak caused by the ECSA lineage in June 2016 in Maceió, Alagoas states, northeast Brazil (**Figure 1a**) [34]. Dataset 2 (ECSA-Br) included only the 29 Brazilian genome sequences. Using a robust nonparametric test [35], no evidence of recombination was found in both datasets.

### Maximum likelihood analysis and temporal signal estimation

Maximum likelihood (ML) phylogenetic analyses were performed for each dataset using RAxML v8 [36]. We used a GTR nucleotide substitution model with 4 gamma categories (GTR+4Γ). In order to investigate the evolutionary temporal signal in each dataset, we regressed root-to-tip genetic distances against sample collection dates using TempEst [37]. For both datasets we obtained a strong linear correlation (dataset 1: r^2^=0.93; dataset 2: r^2^=0.84) suggesting these alignments contain sufficient temporal information to justify a molecular clock approach. However, for dataset 1, the Angola/M2022/1962 strain was positioned substantially above the regression line. Previous investigations have suggested this strain may have been the result of contamination or high passage in cell culture [8], so this sequence was removed from subsequent analyses.

### Molecular clock phylogenetic analysis

To estimate time-calibrated phylogenies we used the BEAST v.1.10.1 software package [38]. To infer historical trends in effective population size from the genealogy we used several different coalescent models. Because preliminary analysis indicated oscillations in epidemic size through time (as also expected from national case report data), we used three flexible, non-parametric models: a) the standard Bayesian skyline plot (BSP; 10 groups) [39], b) the Bayesian skyride plot [40], and c) the Bayesian skygrid model [41], with 45 grid points equally spaced between the estimated TMRCA of the CHIKV-ECSA genotype in Brazil and the date of the earliest available isolate, collected in 18 March 2017 [41]. For comparison, we also used a constant population size coalescent model. We tested two molecular clock models: a) the strict molecular clock model, which assumes a single rate across all phylogeny branches, and b) the more flexible uncorrelated relaxed molecular clock model with a lognormal rate distribution (UCLN) [42]. Because the marginal posterior distribution of the coefficient of variation of the UCLN model did not exclude zero (most likely due to the small alignment size), we used a strict molecular model in all analyses. For each coalescent model, Markov Chain Monte Carlo analyses were run in duplicate for 10 million steps using a ML starting tree, and the GTR+4Γ codon partition (CP)1+2,3 model [42].

### Epidemiological analysis

The epidemic basic reproductive number (R_0_) was estimated from monthly confirmed cases, as previously described [31, 43]. Because (*i*) the Asian genotype was circulating in the north region of Brazil since 2014 [1], and (*ii*) we observed a relatively small number of cases both in the notified and confirmed time series, we assume cases from June 2014 and December 2016 did not represent autochthonous transmission of CHIVK-ECSA. We assume a mean generation time of 14 days, as previously reported elswehere [44]. We report R_0_ estimates for different values of the generation time (g) parameter, along with corresponding estimates of the epidemic exponential growth rate, per month (r).

### Web search query data

Available in near-real time, disease-related Internet search activity has been shown to track disease activity (a) in seasonal mosquito-borne disease outbreaks, such as those caused by dengue [42, 82], and (b) in unexpected and emerging mosquito-borne disease outbreaks such as the 2015-2016 Latin American Zika outbreak [45]. Here, we investigated whether we could find a meaningful relationship between Internet search activity and the local chikungunya outbreak in Roraima. Indeed, novel Internet-based data sources have the potential to complement traditional surveillance by capturing early increases in disease-related search activity that may signal an increase in the public’s perception of a given public health threat and may additionally capture underlying increases in disease activity. Internet searches may be particularly important and indicative of changes in disease transmission early during an outbreak, when ongoing information on the virus transmission is obfuscated by a lack of medical surveillance. In addition, Internet search trends may also help track disease activity in populations that may not seek formal medical care. We used the Google Trends (GT) tool [45] to compile the monthly fraction of online searches for the term “Chikungunya”, that originated from Boa Vista municipality (Roraima state), between January 2014 and July 2018. For comparison, GT search activity for the term “Chikungunya” was collected for the same time period for Manaus municipality (Amazonas state). The synchronicity of GT time series and notified and confirmed case counts from Boa Vista and Manaus was assessed using the Spearman’s rank correlation test in the R software [46].

### Data availability

XML files and datasets analysed in this study are available in the GitHub repository (http://github.com/arbospread). New sequences have been deposited in GenBank under accession numbers MK121891-MK121908 (CHIKV-ECSA) and MK134712-MK134713 (CHIKV-Asian).

## Results

Although most CHIKV notified cases in Brazil were reported in 2016 (**Figure 1**), in Roraima, the majority of notified and confirmed cases in Roraima state were reported in 2017 (5,027 notified cases and 3,720 laboratory-confirmed infections). The number of cases in Roraima started increasing exponentially in January 2017, and the outbreak peaked in July 2017.

We attempted on-site portable nanopore sequencing of isolates collected during the early phase of the outbreak (February to March 2017). We selected 15 RT-qPCR+ virus isolates from autochthonous cases in Roraima state (11 from Boa Vista, 1 from Bonfim, and 1 from Iracema municipalities) (**Table 1**) with a cycle threshold (Ct) ≤30 (mean 20.3, range 13.7 – 27.41). We included two isolates from two infected travellers returning to Roraima in December 2014, and an additional five isolates from Amazonas state (all from Manaus municipality), sampled between July 2015 and March 2017. In less than 48 hours genome sequence data was obtained for all selected isolates and in less than 72 hours preliminary results were shared with local public health officials and the Brazilian Ministry of Health. A mean genome coverage of 86% (20x) per base pair was obtained for the sequenced data; mean coverage increased to 90% when focusing on samples with Ct<26 (**Figure 2a**). Coverage of individual sequences and epidemiological information for each sequenced isolate can be found in **Table 1**.

**Fig. 2.**
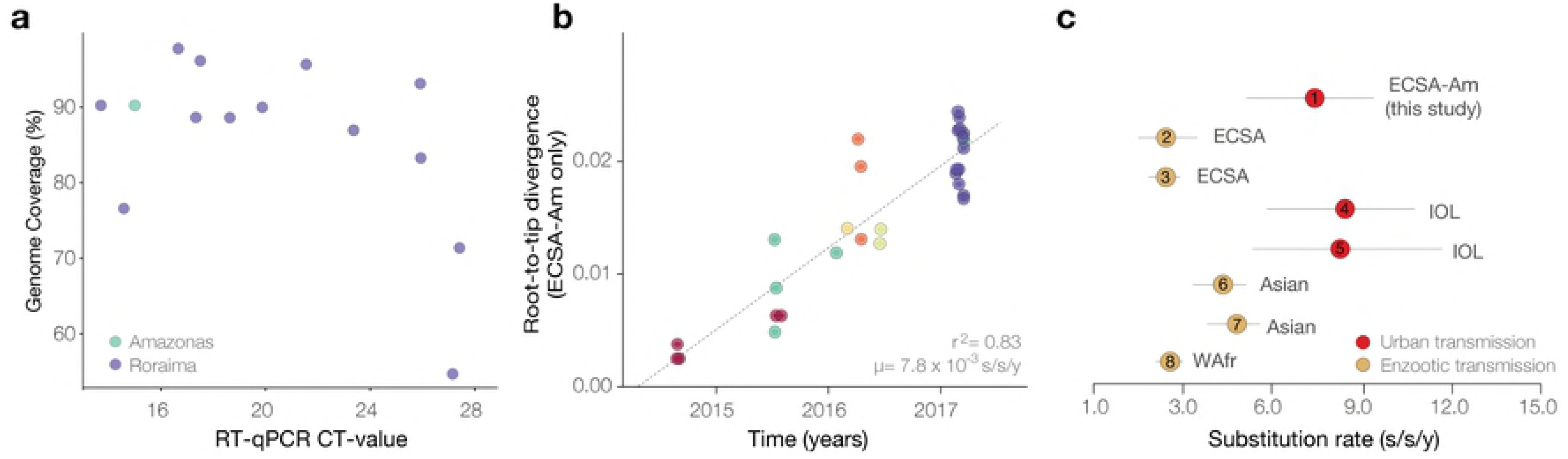
Sequencing statistics, temporal signal and evolutionary rates of the CHIKV-ECSA lineage. A. Genome coverage plotted against RT-qPCR CT-values for the newly generated sequence data. B. Genetic divergence regressed against dates of sample collection for dataset 2 (CHIKV-ECSA-Br lineage). C. Evolutionary rate estimates for the CHIKV-ECSA-Br lineage obtained by this study (circle number 1) compared to published evolutionary rates obtained for other lineages. Circles numbered 2 to 8 represent point estimates reported in [1, 8, 78]. Horizontal bars represent 95% highest posterior density credible intervals for associated evolutionary rates.

Manual and automated phylogenetic analysis identified the ECSA genotype as the dominant genotype circulating in both Roraima and Manaus between 2015 and 2017. However, two cases from late 2014 returning from Venezuela to Roraima (AMA294 and AMA295) were classified as Asian genotype, the dominant lineage circulating in Latin America. Regression analysis of genetic divergence and sampling dates shows accumulation of temporal signal in the ECSA-Br dataset (r^2^ = 0.84) (**Figure 2b**).

We estimated the evolutionary time-scale of the ECSA-Br lineage using several well-established molecular clock coalescent methods. Our substitution rate estimates indicate that the ECSA-Br lineage is evolving at 7.15 x 10^-4^ substitutions per site per year (s/s/y; 95% Bayesian credible interval: 5.04 – 9.55 x 10^-4^). This estimated rate is higher than that estimated for endemic lineages, and is similar to the evolutionary rates estimated for the epidemic lineage circulating in the Indian Ocean region (**Figure 2c**). A closer inspection of amino acid mutations indicate that the ECSA-Br strains lack both the A226V (E1 protein) and the L210Q (E2 protein) mutations that has been reported to increase virus transmissibility and persistence in *Ae. albopictus* populations in the Indian Ocean [47].

ML and Bayesian phylogenetic analyses reveal that the ECSA sequences from Brazil (hereafter named ECSA-Br lineage) form a single well-supported clade (bootstrap support = 100) (**Figure 3**). This is consistent with the establishment of the ECSA genotype in Brazil following the introduction of a single strain to the Americas [1]. The two isolates collected in late 2014 in Roraima cluster together and fall as expected within the diversity of other Asian genotype sequences from the Americas. Our phylogenetic reconstruction suggests at least five separate introductions of the Asian genotype strain Brazil (**Figure S1**), in contrast to a single introduction of the ECSA genotype followed by onward transmission. Moreover, all 13 ECSA isolates sampled in Roraima (*node C*) cluster together with maximum phylogenetic support (bootstrap support = 100; posterior probability = 1.00) (**Figure 3**). We consistently estimate the date of the most recent common ancestor of ECSA-Br Roraima clade to be mid-July 2016 (95% BCI: late March to late October 2016) (**Figure 3**); similar dating estimates under different coalescent models (**Figure S2**). In contrast to the Roraima strains, sequences from Manaus were found to be interspersed with isolates from Bahia and Pernambuco (**Figure 3**), indicating separate introductions of the CHIKV-ECSA lineage, some in early 2015 (*node B*), possibly from the northeast region of Brazil. Interestingly, according to travel history reports, the first autochthonous transmission of CHIKV in Manaus was linked to an index patient who reported spending holidays in Feira de Santana (Bahia state) in early 2015, during a period when this city was experiencing a large CHIKV outbreak [5]. The date of *node A* was estimated to be around mid-July 2014 (95% BCI: early Jul – late Aug 2014), shortly after the arrival of the presumed index case in Feira de Santana, Bahia [5]. This is in line with a single introduction to Bahia (*node A*), followed by subsequent waves of transmission across the northeast and southeast regions of Brazil [5, 48, 49]. Our demographic reconstructions indicate that the outbreak in Roraima 2017 probably represents the third epidemic wave spreading across Brazil (**Figure S3**).

**Figure 3.**
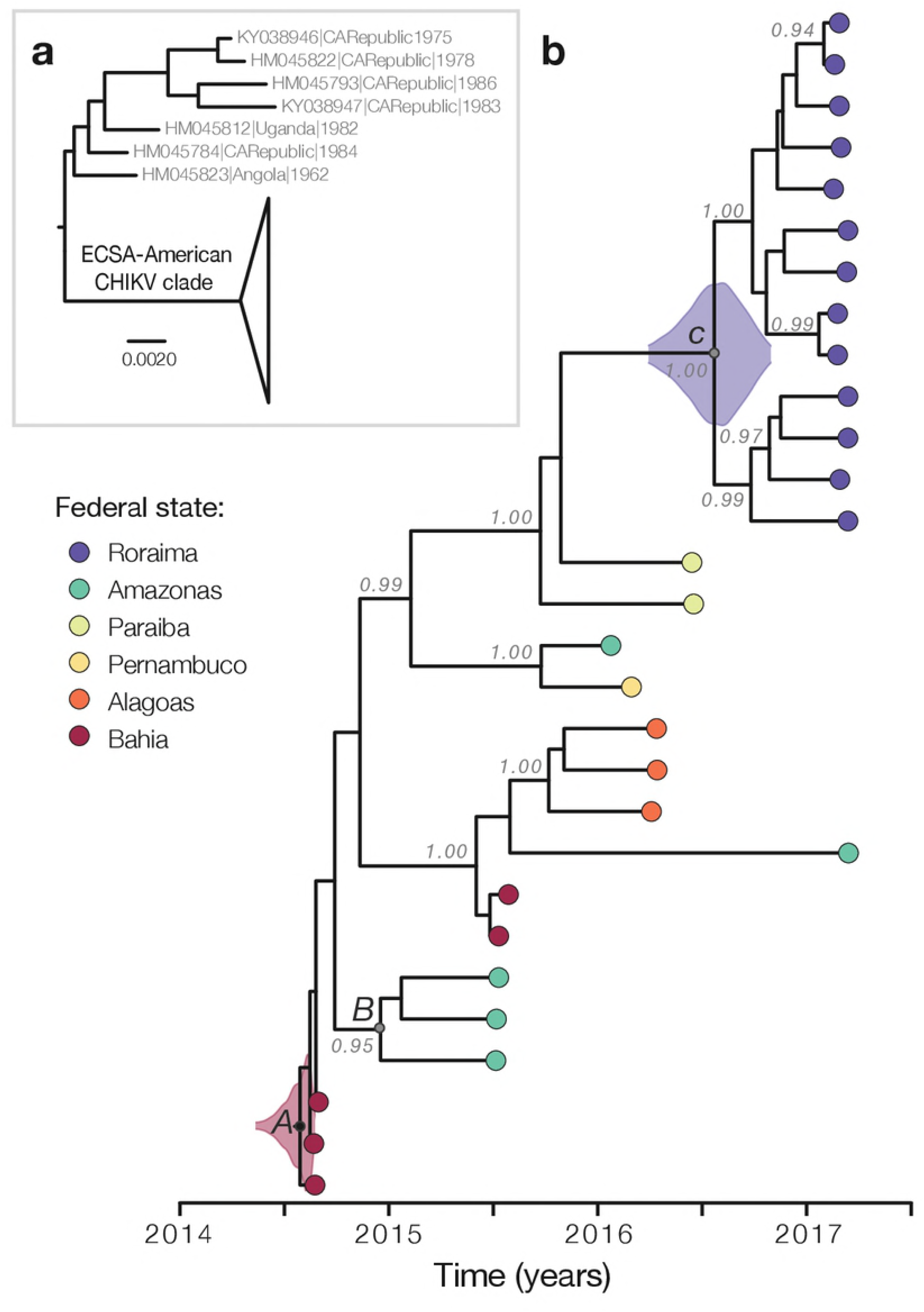
Genetic analysis of the CHIKV-ECSA genotype. A. Maximum likelihood phylogeny depicting the monophyletic clade containing all the Brazilian ECSA isolates (ECSA-Br lineage). B. Time-calibrated phylogeny of all available CHIKV-ECSA whole genome sequences from Brazil, including 18 novel genomes from Roraima and Amazonas states. Colours correspond to state of sample collection. Violin plots show 95% Bayesian credible intervals for associated node heights [38].

Next, we used notified case counts to estimate the basic reproductive number, R_0_, of the epidemic. R_0_ is the average number of secondary cases caused by an infected individual and can be estimated from epidemic growth rates during its early exponential phase [43]. We find that R_0_ ∼1.66 (95% CI: 1.51 – 1.83), in line with previous reports from other settings [50-52]. A sensitivity analysis considering different exponential growth phase periods resulted in a lower bound for R_0_ of around 1.23 (**Figure S4**). To gain insights into the possible magnitude of the outbreak and local surveillance capacity we used the equilibrium end state of a simple susceptible-infected-recovered (SIR) model: *N* = *S* + *I* + *R, S* ∼ 1/R_0_, I ∼ 0, with *N* being the total population size of Roraima. Using this simple mathematical approach, we obtain an attack rate (R) of 0.39 (95% CI: 0.36 – 0.45), slightly lower than elsewhere in Brazil [12, 15]. This corresponds to an estimated 110,882 (95% CI: 102,352 – 127,940) infected individuals, and a case detection rate of 5.34% (95% CI: 4.63 – 5.79). This implies that approximately 1 case was notified for every 19 infections. If we assume 32.7 – 41.2% of the estimated infections are symptomatic, as previously reported in Bahia and Sergipe [53], then we estimate that the local observation success of symptomatic cases was between 12.8 – 16.1%. However, if we assume that 75 – 97% of people infected with CHIKV will develop symptomatic infections, as reported for the Indian Ocean lineage [10, 54, 55], then the chances of reported a symptomatic CHIKV case decrease to 5 – 7% [9]. Case reports suggest that the beginning of the exponential phase of the outbreak was in December 2016 (**Figure S4**), while genetic data suggests that the outbreak clade emerged around July 2016. However, between August 2014 and June 2016, 612 CHIKV notified cases and 40 confirmed cases were reported by the LACEN-RR. It is therefore likely that prior to Jan 2017, low but non-neglectable transmission of the Asian genotype occurred in Roraima.

We investigated the public’s awareness of the chikungunya outbreak by retrospectively monitoring Google searches of the search term “chikungunya” in Roraima state from January 2014 to July 2018 (**Figure 4**). As a comparison, we performed a similar search focusing on the neighbouring state of Amazonas. We found that web search activity and CHIKV cases counts in Roraima are highly correlated (notified cases: *r* = 0.89; confirmed cases: *r* = 0.92, **Figure 4d – e**). Additionally, the timing of the peak of Google searches corresponds to that of notified and confirmed cases with a peak in July 2017 (**Figure 4a and c, Figure 4b and f**). It is important to note that web search activity was available weeks or months before the final number of confirmed (and suspected) cases were made publicly available. This fact highlights the potential utility of monitoring disease-related searches during the outbreak. Interestingly, we find some web-search activity in Roraima before June 2016, particularly in September 2014, March 2015 and March 2016 (**Figure 4f**). These patterns are distinct to those in the Amazonas neighbouring state (notified cases: *r* = 0.65; confirmed cases: *r* = 0.15), which shows an early peak in November 2014, soon after the estimated age of *node B* (**Figure 3b**), followed by a peak in February 2016 and another in March 2017 (**Figure 4c**). These multiple peaks in internet search queries are consistent with the timing of at least 3 introductions detected in our phylogenetic analyses (**Figure 3b**), each possibly resulting in small epidemic waves of CHIKV in Manaus and Amazonas states.

**Figure 4.**
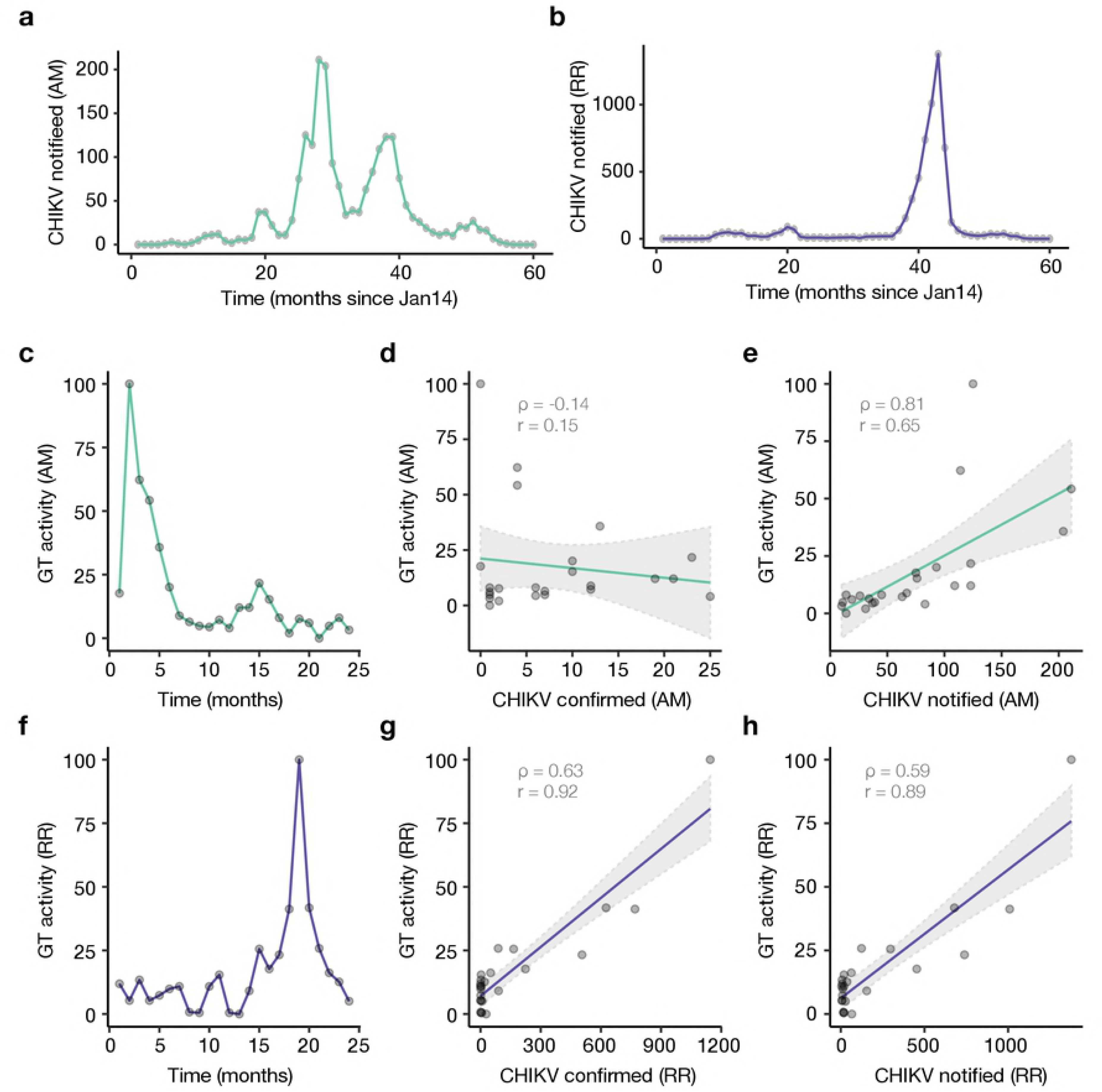
Digital surveillance of chikungunya disease in northern Brazil. A and B show respectively the number of notified CHIKV cases in LACEN-RR and LACEN-AM between Jan 2014 and Sep 2018. Panels C and F show Google Trends activity for the term “chikungunya” in Amazonas (C) and Roraima (F) from Jan 2016 and Sep 2018. Panels D and G show the correlation between Google Trends activity and confirmed cases in Amazonas (D) and Roraima (G), while panels E and H show the correlation between Google Trends activity and notified cases in Amazonas (E) and Roraima (H).

## Discussion

We describe a genomic epidemiological study which used genetic, epidemiological, and digital search data to investigate an outbreak caused by CHIKV in Boa Vista city, Roraima state, northern Brazil, in 2017. Using a combination of genetic, laboratory-confirmed and -suspected, and digital search data from 2014 to 2018, we find evidence for the replacement of the Asian lineage by the ECSA lineage in the north of Brazil. Moreover, we find that ECSA lineage was introduced in Roraima around July 2016, six months before the beginning of the exponential increase in case numbers. Using simple epidemiological modes, we find that on average 1 in 17 (95% CI: 14 – 20) symptomatic CHIKV cases, a fraction of the 110,882 (95% CI: 102,352 – 127,940) estimated number of infections, sought medical care during the outbreak of CHIK ECSA in Roraima. Finally, we find that Google search activity date shows a strong association with CHIKV notified cases in Roraima. Moreover, although the nanopore-based sequencing protocol for CHIKV utilized in this study has been described and validated previously [32], this study represent to our knowledge the first effort to generate on-site complete CHIKV genome sequences. Our results provide evidence of lineage replacement in Brazil, and to the best of our knowledge, deliver a description of the largest outbreak ever reported in north Brazil, revealing the circulation of the ECSA lineage in the Amazon region.

We estimate that 39% (95% CI: 36 – 45)% of Roraima’s population was infected with CHIKV-ECSA-Br during the outbreak in 2017. Our estimates are higher than the 20% seropositive observed in a rural community in Bahia [10], and slightly lower than the 45.7 – 57.1% observed in two serosurveys conducted in the same state [12], where the ECSA lineage also seems to predominate. The observed differences in terms of the proportion of the population exposed to CHIKV in Roraima compared to previous estimates from the northeast region could result from partial protection resulting from low-level transmission of the CHIKV-Asian genotype during 2014 – 2016 in the north region. Alternatively, some level of cross-protection could have been conferred by previous exposure to Mayaro virus (MAYV); Mayaro is an antigenically-related alphavirus that may provide some level of cross-reactivity [56, 57] and is associated with *Haemagogus* spp. vectors [58], but has also been identified in *Culex quinquefasciatus* and *Aedes aegypti* mosquitoes [59]. MAYV has been detected in the north [60-64] and centre-west [21, 59, 65-68] regions of Brazil. Moderate to high prevalence of MYV IgM have been found in urban northern areas [60], which could explain the limited spread of CHIKV in Manaus compared to Roraima.

Different CHIKV circulating lineages may have remarkably different public health consequences. Lineage-specific clinical presentations have been recently highlighted by a recent index cluster study which showed that 82% of CHIKV infections caused by the ECSA lineage are symptomatic, in comparison to only 52% of symptomatic infections caused by the Asian genotype [54]. While the Asian lineage seems to have circulated cryptically for 9 months before its first detection in the Caribbean [3], the faster detection of the ECSA lineage in Brazil could at least in part be a consequence of a higher rate of symptomatic to asymptomatic infections of the ECSA lineage circulating in Brazil. The time lag between the phylogenetic estimate of the date of introduction of a virus lineage and the date of the first confirmed case in a given region, enables us to identify surveillance gaps between the arrival and discovery of a virus in that region [69].

We used genomic data collected over a 3-year period to estimate the genetic history of the CHIKV-ECSA-Br lineage. We estimate that the CHIKV-ECSA-Br lineage arrived in Roraima around July 2016, whilst the first confirmed CHIKV cases in Roraima occurred earlier, in August 2014. That the discovery date anticipates the estimated date of introduction can be explained by initial introduction(s) of the Asian linage (from the north of Brazil or from other south American regions) resulting in only limited onwards transmission, followed by the replacement of the Asian lineages by an epidemiological successful ECSA lineage. Transmission of the Asian genotype during this period is in line with an increase in notified and confirmed cases, as well internet search query data between August 2014 and June 2016. Nationwide molecular and seroprevalence studies combined with epidemiological modelling [70] will help to determine the proportion of cases caused by the ECSA compared to the Asian lineage in different geographic settings, and to identify which populations are still at risk of infection in Brazil.

We estimated high rates of nucleotide substitution for this lineage, which equates to around 8 (95% BCI: 6 – 11) nucleotide substitutions per year across the virus genome. Such rates are similar to the evolutionary rates estimated for the IOL lineage; these are typical of urban and epidemic transmission cycles in locations with an abundance of suitable hosts and lack of herd immunity [8]. None of the mutations associated previously with increased transmissibility of the IOL lineage in *Ae. albopictus* mosquitos in the Indian Ocean region were identified in this study. However, it is currently unclear whether we should expect the same mutations to be linked with increased transmission in *Aedes* spp. populations both from Brazil and from Southeast Asia. Further, it is possible that CHIKV in Brazil is vectored mainly by the *Ae. aegypti* vector that is abundant throughout Brazil [71]. In line with this, CHIKV-ECSA was recently detected in *Aedes aegypti* from Maranhão [72] and Rio de Janeiro states [73].

The past dengue serotype 4 genotype II outbreak in Brazil ignited in the north of the country, and is inferred to have been introduced from Venezuela to Roraima, before spreading to the northeast and southeast region of Brazil [74]. Our genetic analysis reveals at least four instances of ECSA-Br virus lineage migration in the opposite direction, i.e., from northeastern to northern Brazil. Such a pattern may not be surprising due to the year-round persistence of *Aedes aegypti* mosquitos in the northeast and the north areas [31]. Within-country transmission will be dictated by human mobility, climatic synchrony, and levels of population immunity. Moreover, international spread of the ECSA-Br linage is expected to regions linked to Brazil. Previous analyses of dengue virus serotypes has identified a strong connectivity between north Brazil and Venezuela [25, 75], and northeast Brazil and Haiti [31, 76]. In addition, Angola and Brazil are linked by human mobility and synchronous climates that have facilitated the migration of CHIKV-ECSA [1] and Zika virus (http://virological.org/t/circulation-of-the-asian-lineage-zika-virus-in-angola/248).

Improving surveillance in the Amazon region may help anticipate transmission of vector-borne diseases and also spillover from wild mammals of zoonotic viruses of particular concern [27]. Genomic portable sequencing of vector-borne viral infections in the Amazon may is particularly important in the context of early identification of circulation of strains newly (re)-introduced from wildlife. For example, yellow fever strains collected in Roraima seem to be at the source of the 2016-2018 yellow fever virus outbreak in southeast Brazil, which has affected large urban centres in Minas Gerais, São Paulo and Rio de Janeiro [26]. In the near future, the increasing rapidity and decreasing cost of genome sequencing in poorly sampled areas, combined with emerging theoretical approaches [77], will facilitate the investigation of possible associations between arbovirus lineage diversity, mosquito vectors, reservoir species, and transmission potential.

Finally, the reported synchronicities between notified chikungunya case counts in Roraima and the chikungunya-related Internet searches originated in the region highlight the potential complementarity that Internet search activity may offer in future disease outbreaks. Specifically, given that disease-related search activity can be monitored in near-real time, early signals of increases in disease activity may be spotted weeks or months before lab-confirmed case counts may be available in an unfolding outbreak.

## Acknowledgments

We are thankful to all personnel from SVS/MS, PAHO/Brazil, Roraima and Boa Vista Health Surveillance System that coordinated surveillance and helped with data collection and assembly. We thank Oxford Nanopore Technologies for the support to the ZIBRA-2 (Zika in Brazil Real time Analyses-second round) project with additional flowcells and corresponding reagents, and also thank QIAGEN for donation of consumables.

## Supplementary Figure Legends

**Figure S1. Maximum likelihood phylogenetic tree of the CHIKV Asian genotype.** Includes isolates from Southeast Asia, Americas and Brazil. Isolates represented by blue tips were sampled in Roraima, while isolates shown in red represent other strains sampled in Brazil.

**Figure S2. Dating estimates obtained under different coalescent models.** Estimates for node A (time of the most recent common ancestor, in dark red, see Figure 3b), node B (main Amazonas clade, in green), and node C (Roraima clade, in purple) are shown for different non-parametric models (Bayesian skygrid, skyride, skyline) and for a simple constant population size model.

**Figure S3. Demographic dynamics of CHIKV ECSA-Brerican lineage in Brazil.** Fluctuation of effective population size over time as inferred through a Bayesian skygrid coalescent model.

**Figure S4. Exponential Period of the CHIKV epidemic in Boa Vista municipality, Roraima state.** Log number of notified cases per month are plotted against number of months since January 2015.

## Supplementary Table

**Table S1.**
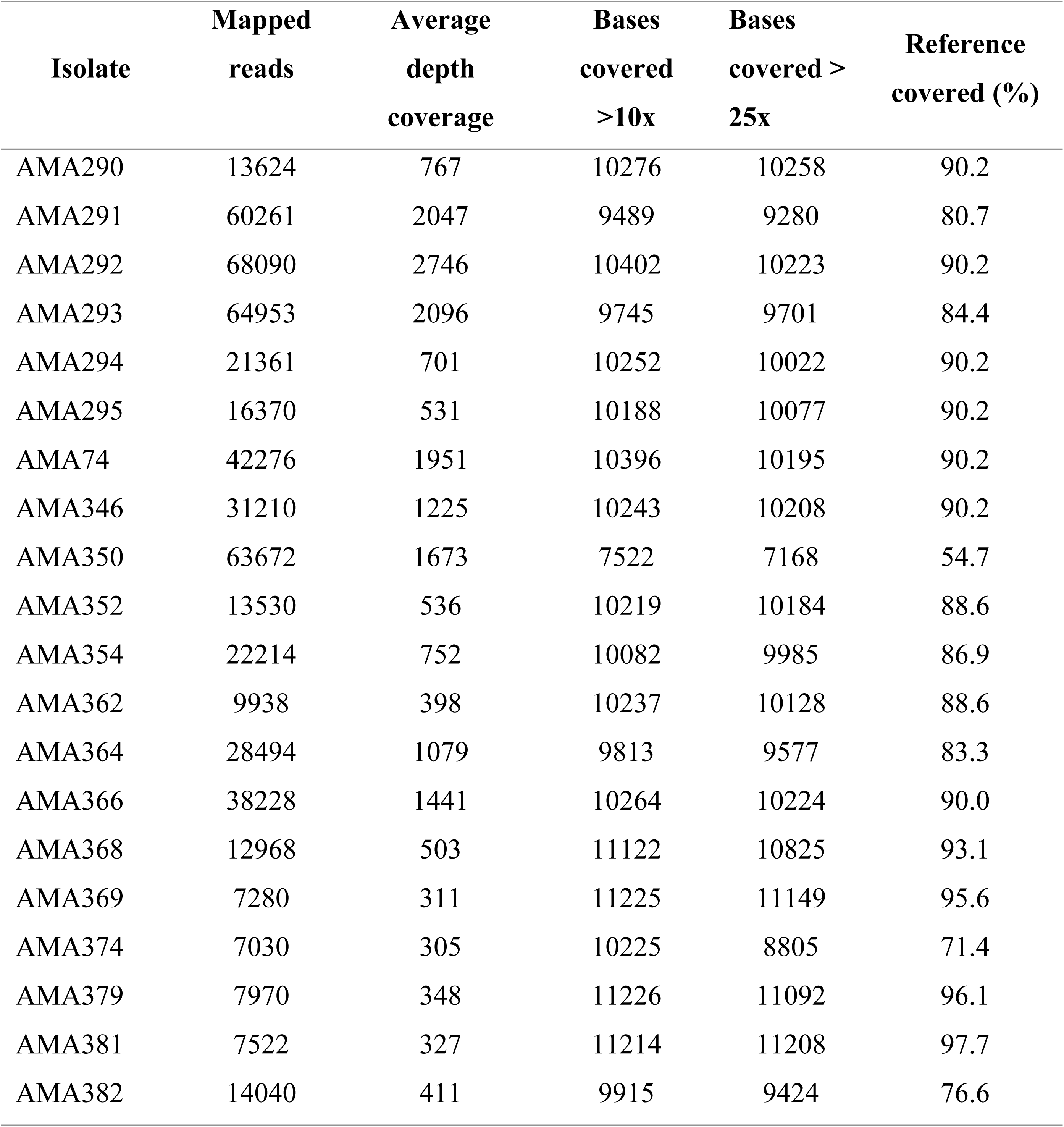
Minion sequencing statistics.

